# RAGE antagonist peptide mitigates AGE-mediated endothelial hyperpermeability and accumulation of glycoxidation products in human ascending aortas and in a murine model of aortic aneurysm

**DOI:** 10.1101/2021.10.22.465199

**Authors:** Chiara Camillo, Alexey Abramov, Philip Allen, Estibaliz Castillero, Emilia Roberts, Yingfei Xue, Antonio Frasca, Vivian Moreno, Mangesh Kurade, Kiera Robinson, David Spiegel, Damien LaPar, Juan B Grau, Richard Assoian, Joseph E Bavaria, Hiroo Takayama, Giovanni Ferrari

## Abstract

**Background:** Aortic dissection and aneurysm are the result of altered biomechanical forces associated with structural weakening of the aortic wall caused by genetic or acquired factors. Current guidelines recommend replacement of the ascending aorta when the diameter is >5.5 cm in tricuspid aortic valve patients. Aortopathies are associated with altered wall stress and stiffness as well as endothelial cell dysfunction and synthetic vascular smooth muscle cell (VSMC) phenotype. We reported that these mechanisms are mediated by glycoxidation products [Reactive oxygen species (ROS) and Advance Glycation End products (AGE)]. This study addresses the role of glycoxidation on endothelial function and AGE-mediated aortic stiffness.

**Hypothesis and aims:** Here we investigate how circulating glycation products infiltrate the aortic wall via AGE-mediated endothelial hyperpermeability and contribute to both VSMC synthetic phenotype and extracellular matrix (ECM) remodeling *in vivo* and *ex vivo*. We also study how RAGE antagonist peptide (RAP) can rescue the effect of AGEs *in vitro* and *in vivo* in eNOS^−/−^ vs WT mice.

**Methods and results:** Human ascending aortas (n=30) were analyzed for AGE, ROS, and ECM markers. *In vitro* glycation was obtained by treating VSMC or human and murine aortas with glyoxal. Endothelial permeability was measured under glycation treatment. Vascular stiffness was measured by a pressure myograph comparing wild-type mice ± glyoxal. eNOS^−/−^ mice, a model of increased endothelial permeability, were treated for 28 days with hyperlipidemic diet ± Angiotensin II (1000ng/kg/min) with or without anti-glycation treatment (RAP 20mg/kg). Echo data of aortic diameter were collected. Murine vascular stiffness was measured by a pressure myograph (n=5/group). Glycoxidation products were detected in all human aortas independently of aortic diameter, with stronger accumulation on the lumen and the adventitia layer. AGEs increased endothelial permeability, induce synthetic phenotypic switch in human VSMCs, and inhibit cell migration. RAP pre-treatment rescue the effect of glyoxal on endothelial cells. Ex vivo glycation treatment of murine arteries impacted on ECM and increased stiffness. Aortic stiffness was higher in eNOS^−/−^ vs WT mice. Ang II-mediated aortopathies results in aortic dilation, and AGE/ROS accumulation, which is rescued by RAGE antagonist peptide treatment of eNOS^−/−^ mice.

**Conclusions:** Glycoxidation reaction mediate EC permeability, VSMCs phenotype, and ECM remodeling leading to dysfunctional microstructure of the ascending aorta, altered vascular stiffness and increasing aortic susceptibility to dilation and rupture. Moreover, we show that RAP can mitigate AGE-mediated endothelial hyper-permeability *in vitro* and impact on ascending aneurysm *in vivo*

## INTRODUCTION

Thoracic aortic aneurysm (TAA) and dissection are major causes of morbidity and mortality in the US. Current American heart association (AHA) guidelines suggest replacement the ascending aorta when the diameter is >5.5 cm in tricuspid aortic valve (TAV) patients^1, 2^. More aggressive surgical repair is recommended for patients with congenital heart defects like Marfan Syndrome or Bicuspid Aortic Valve (BAV) undergoing AV replacement^3^. However, clinical evidence shows that aortic dissection and rupture often occur outside of the suggested metric-based parameters. Aortopathies, like aortic dissection and aneurysm, are the result of altered biomechanical forces associated with structural weakening of the aortic wall caused by genetic and acquired factors^4^. It has been previously reported that a significant fraction of patients with acute type A aortic dissection have aortic diameters lower than 5.5 cm^5, 6^. A major obstacle hampering the decisions of whether to replace the ascending aorta is the absence, in the current guidelines, of tools informing on the integrity and stiffness of the aortic wall independently of metric measurements. Imaging diagnostic methods are imperfect predictors as they do not inform on the structural integrity of the aortic wall structure^5^ and the impact of clinical comorbidities, disease propensity, and individual variations in blood biochemistry are only marginally considered in aortic surveillance programs. Thus, identifying new mechanisms impacting aortic wall integrity could have major significance for the development of novel molecular and imaging biomarkers aimed at risk-stratifying patients with ascending aortic diseases.

Glycation is a well-described contributor to cardiovascular disease, notably in diabetic patients and in aging^7^. Advanced Glycation End-Products (AGEs) are irreversible aggregates that result from a multistep reaction beginning with non-enzymatic protein glycation, or lipid peroxidation^1–3^. A major mechanism through which AGEs contribute to disease arises from structural protein cross-linking in tissue wall (type 1 collagen and elastin) and from their direct impact on cellular phenotype and proliferation/apoptosis^4–6^. This leads to alterations in structure and function of the glycated proteins, and to generation of oxidative stress via the induction of Reactive Oxygen Species (ROS)^6,7^ formation. Central intermediates in the pathway to AGE formation include precursors such as reactive carbonyl species, glyoxal, and methylglyoxal. These key species are the precursors to AGEs, a heterogeneous group of molecules. AGEs could signal through the receptor for advanced glycation end products (RAGE), a cell surface receptor present on cardiac myocytes and blood vessels^8^ leading to oxidative stress via ROS^9^ and/or participate in the inflammatory response. Thus, glycoxidation products contribute to tissue degeneration via two pathways: modulation of cellular phenotypes via multiple inflammation-related receptors—most prominently including RAGE—as well as modification of cytoplasmic proteins, and biochemical modification and concomitant structural/functional degeneration of ECM components such as collagen and elastin crosslinks.

An association between valve and vascular dysfunctions and AGEs has been described recently ^1, 10^, showing the role of RAGE in mediating VSMC contractile-to-synthetic phenotypic switch; this contributes to ECM remodeling by proteinases (i.e. Matrix metalloproteinase (MMPs))^7, 11^. Previous works have emphasized how AGE/RAGE pathway has a role in affecting vascular barrier permeability in diabetes and its complications^12, 13^. In addition, AGEs and glycated circulating serum proteins are known to permeate through the endothelium in the aortic wall. Studies document the formation of glycation products in various blood cells and endothelial cells^76,86^. In fact, while many studies report responsiveness of VSMC to treatment with preformed AGEs, there is no equally strong evidence of AGE formation in vascular media. *In situ* glycation could require infiltration of preformed glycation products or reactive glycation precursors into the aortic wall via impaired endothelial barrier. These glycated proteins could then be crosslinked to large ECM molecules altering collagen and elastin architecture^14^. In recent studies we have reported how glycation and permanent incorporation of circulating proteins, results in collagen misalignment via cross-link formation^8,9^. Histological analysis of human ascending aortas showed that soluble RAGE (sRAGE) correlates with dysfunctional aortic microstructure and does not correlate with aortic diameter or diameter/body surface area^10^. We also described how oxidation reactions modulate VSMC phenotype and correlated with the area of altered wall stress and microstructural dysfunction of the ascending aorta^11^. ROS also results in oxidized aminoacidic accumulation, such as di-tyrosine, a cross linker that could affect ECM stiffening.

Glycoxidation accumulation in the vascular wall correlates with age-related aortic stiffness *in vivo*^15^ and ECM remodeling affects arterial mechanics altering wall stiffness. Moreover, MMPs, AGEs, and angiotensin have been implicated in arterial stiffening, resulting in altered wall stress and endothelial dysfunction^16, 17^. Taken together, these data brought us to formulate the hypothesis that impaired endothelial permeability could lead to pathological accumulation of glycated proteins in the sub-endothelial vascular structure, resulting in phenotypic changes of VSMC and remodeling of ECM. These cellular and matrix alterations, which are independent of aortic diameter, could affect aortic stiffness. Notably, patients with type 2 diabetes mellitus (T2DM) had significantly reduced risks of aortic aneurysm (AA) and aortic dissection (AD). It has been suggested that glycated cross-links in aortic tissue may play a protective role in the progression of aortic diseases among patients with T2DM^18^. Here we aim to understand how AGEs impact endothelial permeability, endothelial cell phenotype, and vascular stiffness using primary cells, *in vitro* glycation, and by using animal models with known endothelial dysfunction. Furthermore, we investigate a high-affinity RAGE-specific antagonist peptide (RAP), FPS-ZM1, that blocks AGEs binding to the V domain of RAGE inhibiting the downstream pathways, as a tool to rescue AGE-mediated endothelial permeability *in vitro* and *in vivo*. This study could help explain the role of glycoxidation crosslinkers accumulation in patients with pathologically-of pharmacologically-enriched circulating AGE products.

## RESULTS

### Glycation products permeate human ascending aortas, independently of aortic diameter, in both the intima and adventitia layer

Thirty patients (21 males and 9 females, age mean 64.11 ± 10.2; aortic diameter from 3.4 to 7 cm) with non-syndromic aortopathies and tricuspid aortic valve morphology undergoing ascending aortic replacement were selected from our aortic biobanks. The characteristics of the study population are shown in **Table S1**. Irrespectively of aortic diameter, expansion rate, and ratio of aortic area/diameter to body weight/surface all resected aortas show some level of elastin and ECM remodeling (**Figure 1A**). To assess the presence of glycation products these specimens were analyzed for generalized AGE staining, carboxymethyl-lysine (CML)^19^, AGE-crosslinker glucosepane, and extracellular Modified Movat pentachrome to study the distribution and integrity of collagen fibers and proteoglycans. Our data revealed an abundant expression of generalized AGE staining in both dilated and non-dilated tissues, regardless the diameter of the ascending aorta, sex, and key co-morbidities (**Figure 1 and Figure S1**). Interestingly, the pattern of staining shows strong accumulation in both the intima (I) and adventitia (A) layers of the aorta (**Figure 1B**). Media layer in larger vessels is practically avascular; thus, nutrients and oxygen diffuse to the media from the lumen of the vessel and from the vessels of the adventitia. Diffusion of solutes to the media is supplemented by vasa vasorum and the nutrients and compounds permeates half or two-third of the media^20, 21^. This is consistent with the pattern of generalized AGE and glucosepane accumulation seen in human aortic tissues stained^45^ (**Figure 1B, *a* and *c***). Many cells have developed intrinsic detoxifying pathways against accumulation of AGEs, thus - as expected from surgically removed tissues - CML, a major product of oxidative modification of glycated proteins, only moderately stain the aortic specimens (**Figure 1B, *b***). However, CML can be generated *in vitro* by a reactive sugar aldehydes, glyoxal^22^. In order to implement a model of *in vitro* glycation to study the mechanistic impact of AGEs in endothelial integrity and function, we treated human aortic tissue with increased concentration of glyoxal (500 μM to 50 mM). Compared to the untreated sample (**Figure 1C, *a***), we observed an accumulation of CML at higher doses of glyoxal (**Figure 1C, *b* and *c***) that also correspond to areas of increased elastin degradation, represented by EVG staining (**Figure 1C, *d* and *f***), mimicking areas of ECM dysfunction seen in diseased aortas (**Figure 1A, *a* and *b***). Having shown that AGEs are detectable in human aortic specimens and can be modulated *in vitro*, we aimed to assess their role on the main cellular and ECM components of the ascending aortas.

**Figure 1:**
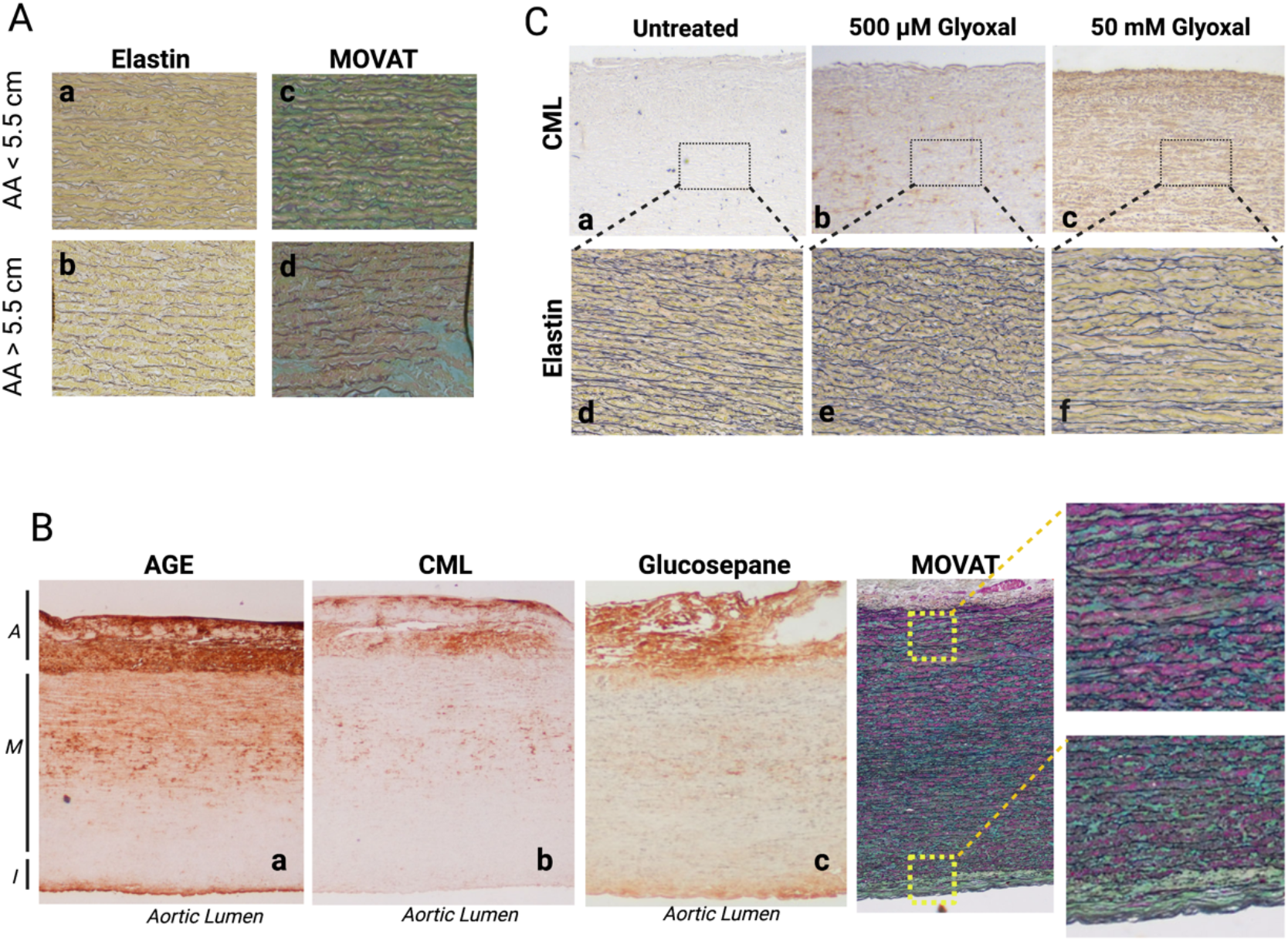
Glycation products are positively stained in aortas, independently of aortic diameter. Human ascending aorta tissues collected from TAV/TAA patients. **(A)** Paraffin fixed aortic tissues have been stained with Van Gieson’s or Movat staining for ECM components. IHC staining performed with an anti-AGE or anti-CML antibody for glycation compounds in ascending aorta tissue from TAA and non-dilated patients. Upper panels (***a, c***): representative section of a TAV AA 5.5 cm < diameter ascending aortic tissue. Lower panels (***b, d***): representative section of a TAV AA 5.5 cm > diameter ascending aortic tissue. Images are representative of staining performed on n=5 tissues/group of patients; 20x magnification. **(B)** Human ascending aorta cross-section showing the three layers of the aortic wall: adventitia (A), media (M), intima (I). IHC of AGE, CML, glucosepane and Movat staining. **(C)***In vitro* induced glycation. Fresh ascending aorta has been treated or not with increased doses of DMEM 10% glyoxal (500μM and 50mM glyoxal) at 37°C for 24h. IHC staining for CML for glycation compounds, or EVG staining for elastin have been performed on paraffin fixed aortic tissue biopsy punches.

### RAGE antagonist peptide rescues AGE-mediated endothelial phenotype and permeability

Studies document the formation of glycation products in various blood cells and endothelial cells. *In situ* glycation requires infiltration of preformed glycation products and/or reactive glycation precursors into the aortic wall via impaired endothelial barrier. Results shown in **Figure 1** support this idea. Thus, we designed *in vitro* assays aimed at investigating the role of glycation precursor compound on aortic endothelial cells. First, we performed an *in vitro* collagen matrix-contracting assay on bovine-derived aortic endothelial cells with results showing that glyoxal significantly impaired the ability of ECs to contract the collagen matrix after 24h of treatment (**Figure 2A**). Glyoxal also decreased the migration rate of ECs significantly affecting the wound-healing closure (**Figure 2B**). To further investigate the biological roles of AGEs on aortic endothelium, we studied the *in vitro* effect of glycation on endothelial permeability by utilizing the Z-Theta ECIS system with the purpose of monitoring the barrier function, defined as resistance (R), of a cell monolayer. After the formation of a cell monolayer given by the resistance (ohm) value plateauing, ECs were stimulated with or without glyoxal and monitored over time. ECs treated with glyoxal showed a significant decrease of resistance (increased permeability) compared to not treated (NT) cells several hours after treatment. AGEs effect on endothelial monolayer permeability has been previously described^12, 23^. However, whether this effect is mediated by RAGE has not previously been addressed and in our experiment the treatment with RAP (FPS-ZM1 5 and 10 μg/ml) effectively rescued the increased permeability to control levels in ECs (**Figure 2C**).

**Figure 2.**
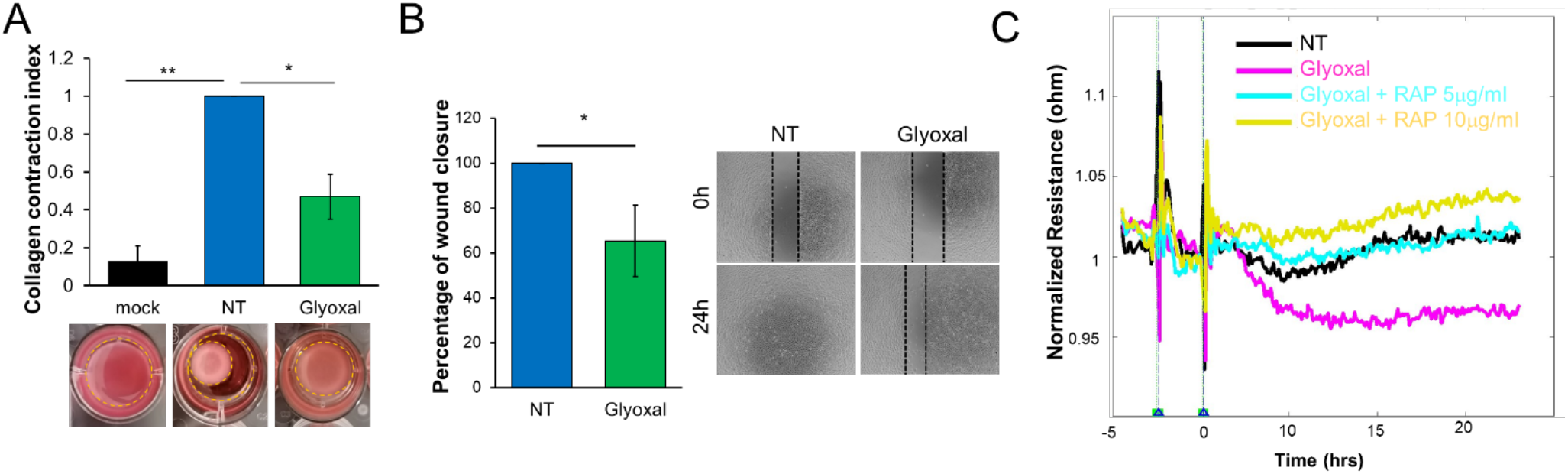
*In vitro* glycation impacts on endothelial phenotype and permeability. (**A**) Images and quantification of collagen matrix contraction assay. Representative images of a collagen disk without cells (mock), PBS (NT) or glyoxal treated (500μM 24h) of BAOEC cells. Results are the average ± SEM of 4 independent assays. Statistical analysis: one-way ANOVA Bonferroni postdoc analysis; p≤0,05 *, p<0,01 **. **(B)** Comparison between ECs treated or not with glyoxal (500μM 24h) migration rate in wound healing assays expressed as percentage of wound closure compared to the control. Results are the average ± SEM of 3 independent assays. Data are normalized on T0. Statistical analysis: t-test; p≤0,05 *. **(C)** Endothelial monolayer permeability was assessed by ECIS Zeta-Theta. When ECs reach the monolayer (~ 48h), cells were treated or not (black) with RAP [5μg/ml] (blue) or [10μg/ml] (yellow) for 2h and then with glyoxal 500μM (pink). The effect of the treatment on the Resistance (R, ohm) value has been monitored during time. R values are normalized at a precise time point, immediately before adding the treatment on the cells. Wells without cells has been used as blank (internal control) for the experiment. A representative graph of the normalized resistance values of 4 independent experiments conducted under identical conditions.

### RAGE antagonist peptide restores VE-cadherin membrane localization impaired by glycation treatment in endothelial cells

Regulated protein phosphorylation and endocytosis mediates the disruption of adherens junctions (AJs) by permeability agents^24^. In particular, several studies reported that variation in the expression of VE-cadherin and its binding partners induced by TNFα, VEGF^25^ or histamine^26^, correlates with the destabilization of endothelial junctions. Considering that VE-cadherin acts downstream of Src^27^ causing the disruption of the endothelial barrier in response to permeability agent, we studied how the regulation of VE-cadherin expression is affected by glycation treatment^28^. As shown in (**Figure 3A-B)**, when compared to not-treated cells, VE-cadherin expression level is decreased after glyoxal treatment significantly and it is partially rescued by a pre-treatment with RAP. To explore the involvement of AGEs in the AJs formation and organization, we plated ECs on gelatin-coated coverglass and assessed, by confocal microscopy, VE-cadherin targeting at cell-to-cell contacts. Immunofluorescence confocal analysis shows how cell-cell VE cadherin localization, expressed in the AJs of ECs, is impaired by glyoxal treatment causing a decrease of membrane VE-cadherin localization (**Figure 3C**). Indeed, VE-cadherin is less expressed in the cell-to-cell contacts and instead accumulated in the cytoplasm after glyoxal treatment. As previously reported, the decreased barrier function could be related to the internalization of VE-cadherin that can explain the increased permeability^28, 29^. RAP treatment prevented the diminished VE cadherin membrane localization caused by glyoxal (**Figure 3C**). These data suggest that glycation not only decreases the ability of the ECs to contract ECM or migrate, but also contributes to increasing VE-cadherin-mediated endothelial permeability.

**Figure 3.**
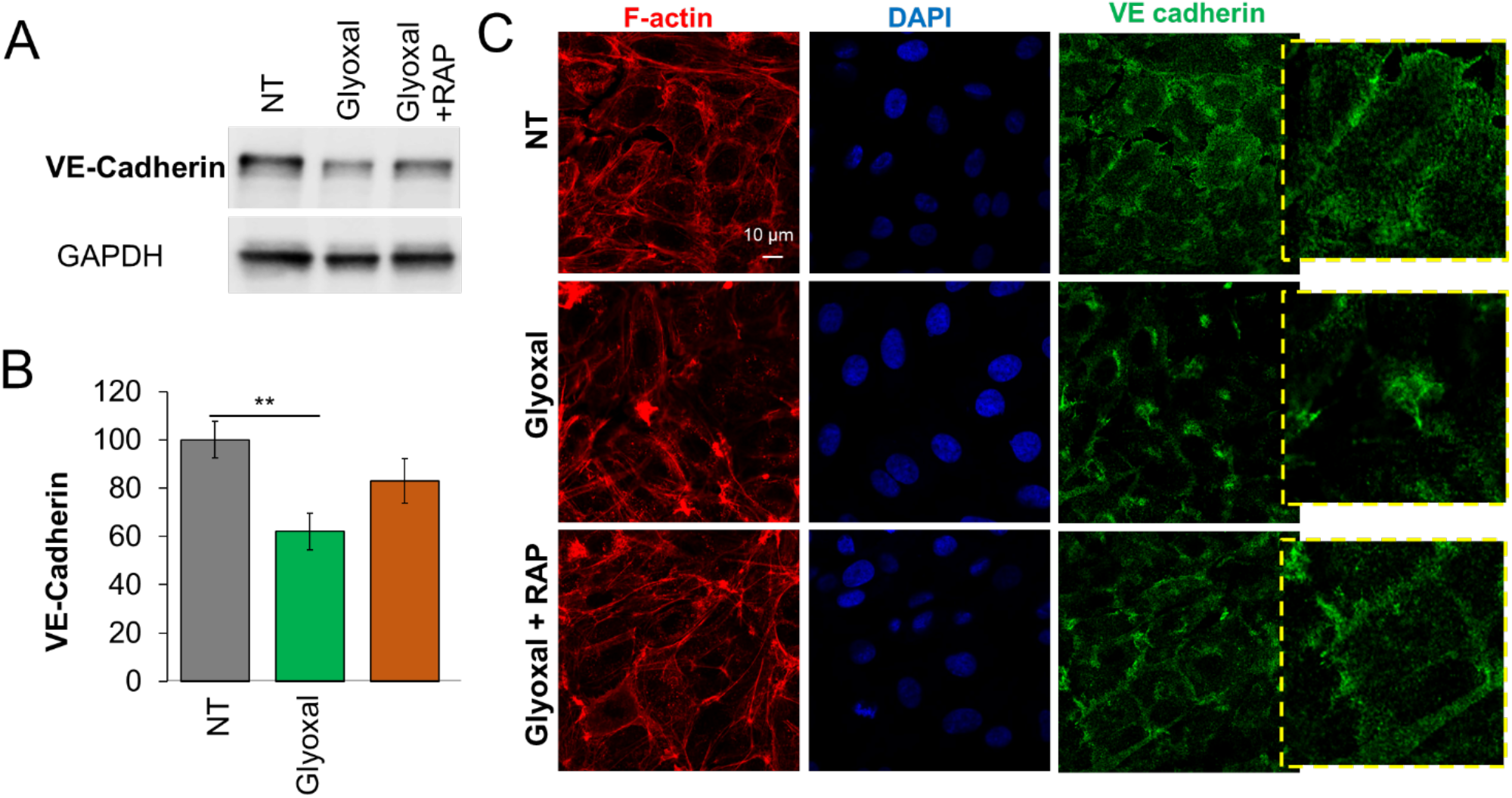
RAGE antagonist peptide restores VE-cadherin membrane localization impaired by glycation treatment in endothelial cells. (**A**) Representative western blot of VE-cadherin protein level in ECs pre-treated with RAP [5μg/ml] for 2h and then with glyoxal (500μM) for 24h. (**B**) Results are the mean ± SEM of n≥ 4 independent experiments (a representative one is shown). Statistical analysis: one-way ANOVA and Tukey’s post hoc analysis. p<0,01 **. (**C**) Fluorescence confocal microscopy reveals that ECs treated with glyoxal 500μM show a decrease in VE-cadherin (green) cell-cell localization compared to NT ECs. A pre-treatment with RAP [5μg/ml] rescues the effect. Scale bar 10 μm.

### *In vitro* glycation influences VSMC phenotype

A more permeable endothelium results in accumulation of glycoxidation products in the media layer of the aorta, as suggested by **Figure 1B**. To determine the impact of glycation on aortic VSMC phenotype in the media of ascending aortas, we exposed human aortic smooth muscle cells (HASMCs) to glyoxal and analyzed the expression of synthetic and contractile phenotypes markers. Previously, the appropriate concentration of glyoxal was established through a viability assay on HASMCs (**Figure S2**). qPCR data show that AGEs decrease significantly the contractile phenotype marker levels, including caldesmon, calponin, myocardin, SMMHC and SMA (**Figure 4A**), while increase the synthetic ones like vimentin, CTGF (**Figure 4B**). As we aimed to determine if glycation impacts on the ability of VSMCs to adhere and contract the ECM, we also conducted an *in vitro* collagen matrix-contracting assay. **Figure 4C** show that glyoxal-treated HASMC ability to contract collagen matrix is reduced compared to untreated cells. Next, we investigated whether and how glycation could influence motility of HASMCs on ECM. As shown in **Figure 4D-E**, glycation treatment results in a significant reduction of the migration rate of HASMCs in closing the wound compared to untreated cells after 24h. Thus, glycation treatment induces the shift of the HASMCs towards synthetic phenotype, decreasing the cell ability to contract the collagen matrix and impairing their migration potential. These data expand previous reported data linking AGE-RAGE pathway to HASMC activation, proliferation and regulation of contractile functions ^30^.

**Figure 4:**
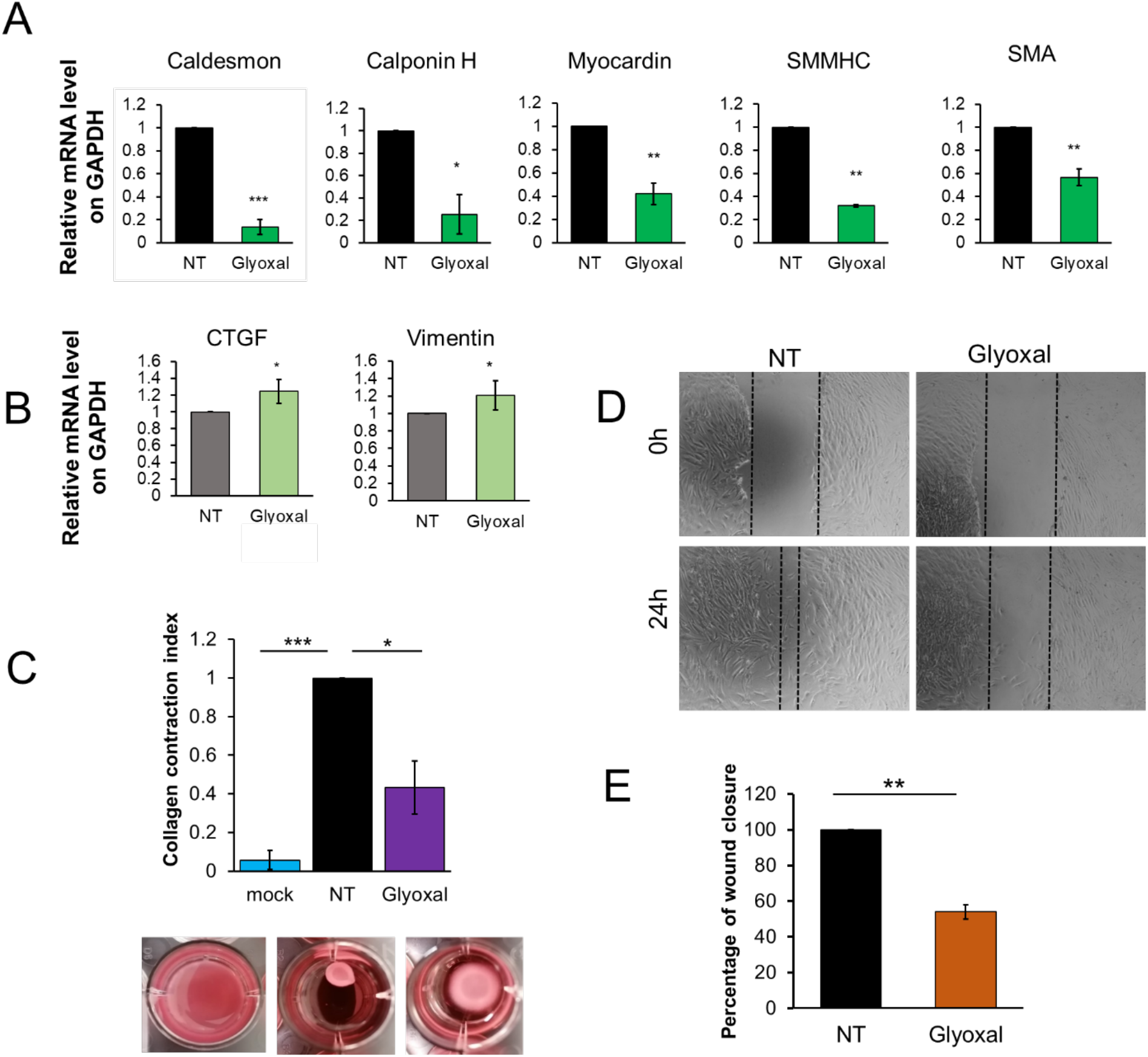
*In vitro* glycation impacts on VSMC phenotype. Real time quantitative PCR analysis of HASMCs treated or not with 1mM glyoxal for 24h. Markers of both **(A)** contractile (caldesmon, calponin H, myocardin, SMMHC, SMA) or **(B)** synthetic (CTGF and vimentin) were measured. NT = untreated. Results are the average ± SEM of ≥ 3 independent experiments. Results were analyzed by a two-tailed heteroscedastic Student’s t-test. **(C)** Images and quantification of collagen matrix contraction assay. Representative images of a HASMCs collagen disk; mock (without cells), NT (PBS) or glyoxal treated (1mM, 24h). Results are the average ± SEM of ≥ 3 independent experiments. Results were analyzed by a one-way ANOVA Bonferroni postdoc analysis. **(D, E)** Comparison between HASMCs treated or not with glyoxal (1mM, 24h); migration rate in wound healing assays expressed as percentage of wound closure compared to the control (NT) and normalized at T0h. Results are the average ± SEM of ≥ 3 independent experiments. Results were analyzed by a two-tailed heteroscedastic Student’s t-test; p< 0,05 *; p≤ 0,01 **; p≤ 0,001 ***.

### Role of AGE on ascending aortic stiffness: *Ex vivo* glycation greatly increase vascular stiffness in murine arteries

Vascular remodeling implies alterations in the synthesis and degradation of ECM components, particularly elastin and collagen, causing vascular disfunction^31^. To further investigate the role of the AGE signaling *in vivo* and determine its impact on aortic wall stability and ECM remodeling in the presence of altered endothelial function we utilized an established endothelial nitric oxide synthase (eNOS) knock-out murine model. eNOS^−/−^ mice show a severe vascular phenotype characterized by increased endothelial permeability, hypertension, deficiency in angiogenesis, decreased cell sprouting and wound repair^31, 32^. A pressure myography assay was used to determine the stiffness of the vessel as per our previous works^16 12^. We performed biaxial inflation-extension tests on C57BL/6 and eNOS^−/−^ mice to study the effects of a vascular deficient-phenotype mouse model on arterial biomechanics. A DMT model 114P Pressure Myograph was used to determine the stiffness of the vessels (**Figure S3A**). Unloaded vessel outer diameter (**Figure S3B and C**) and inner radius (**Figure S3D**) vary significantly with the deletion of eNOS, while wall thickness does not (**Figure S3E**). These data show that arteries become less compliant in a vascular deficient phenotype tissue. We then used pressure myography to study the effect of eNOS knock out response to physiological arterial pressure axially and circumferentially. Interestingly, the compliance defect was more evident in the derived circumferential stress-stretch relationship, which showed a statistically significant leftward shift of the stress-stretch curves in eNOS^−/−^ mice compared to WT especially within physiological pressure range (black symbols, 80-120 mmHg) (**Figure 5A**). Further, the intersection of force-stretch curves defined *in vivo* stretch (IVS) (**Figure S3F**) and the axial stress-stretch curves at 100mmHg showed no relevant difference between WT and eNOS^−/−^ arterial stiffening (**Figure S3G**). These data show that eNOS^−/−^ phenotype results in increased circumferential stiffening of the arteries when compared to wt. We previously described a relevant *in vitro* glycation assay with the goal of increasing the formation of AGEs after treatment with glyoxal in human aorta (**Figure 1C**) and pericardial tissues^33^. Thus, murine arteries from WT animals were exposed to glyoxal (500μM to 50mM) or saline. Outer vessel diameter (**Figure S3H**) and inner radius (**Figure S3I**), but not wall thickness (**Figure S3L**) results show a significant difference between saline and high doses of glyoxal. This effect is more apparent in the derived circumferential stress-stretch relationship, which showed a clear leftward shift of the stress-stretch curves with increased glyoxal dose (**Figure 5B**). Difference has been observed in the IVS among the treatment groups (**Figure S3M**). Interestingly, also axial stress-stretch curves at 100 mmHg revealed a significant glyoxal treatment-dependent arterial stiffening, as indicated by the increased stress with high glyoxal dose compared to saline (**Figure S3L**). Hence, treatment with glyoxal in vitro is sufficient to demonstrate the effect of a vascular-deficient model in aortic stiffness. Moreover, to strengthen our hypothesis that the *in vitro* glycation induced-stiffening was due to an ECM remodeling, we first analyzed the elastin component of the tissues through second-harmonic generation (SHG) microscopy and EVG staining (**Figure 5C**). Control arteries showed organized elastin fibers with both SHG and EVG staining. Compared to saline (**Figure 5C, *a-a’***), instead, glyoxal 50 mM treated arteries were characterized by a severely impacted organization of elastin fibers in both autofluorescence TPEF images (**Figure 5C, *c-c’***) and EVG staining (**Figure 5C, *f***). However, elastin fibers from an explanted carotid of a glyoxal 500μM treated mice were not affected by the treatment as both SHG images (**Figure 5C, *b-b’***) and Verhoeff’s staining revealed (**Figure 5C, *e***). We also observed collagen fibers morphology through the SHG imaging analysis, and no differences have been noticed between the samples (data not shown). To verify AGE accumulation in the arterial tissue due to *in vitro* glycation treatment, we performed IHC analysis on mice carotids used in the stiffness assay (**Figure 5D**). Interestingly, both CML and AGE staining are increased at a high dose of glyoxal (50mM) highlighting endothelium and adventitia layer of the artery (**Figure 5D, *c* *and f***) compared to the control (**Figure 5D, a *and d***). AGE accumulation is mildly evident with the glyoxal 500μM dose, a concentration that preserves cell viability for both endothelium and VSMC cell layer (**Figure 5D, *b* *and e***).

**Figure 5:**
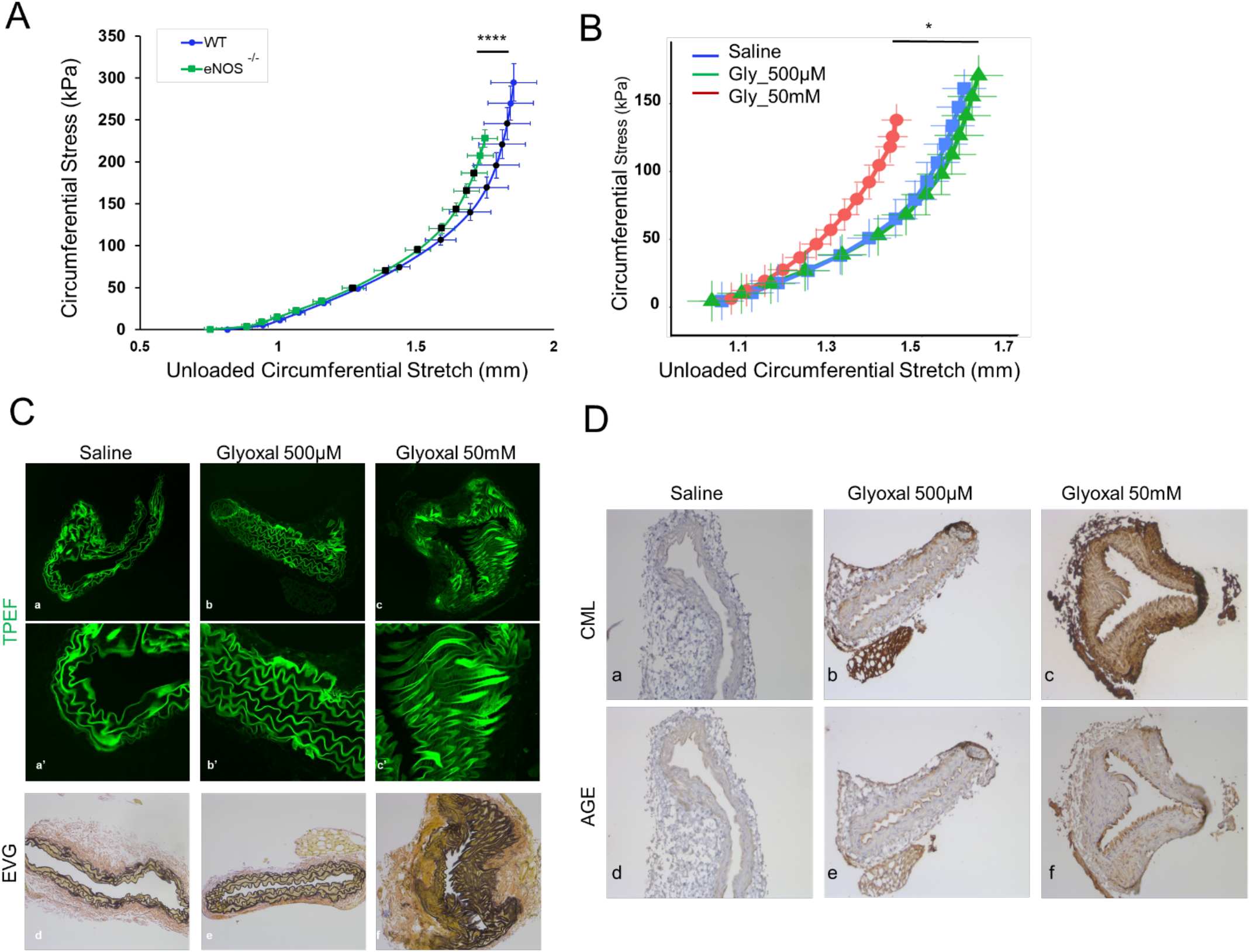
Ex vivo glycation greatly increase vascular stiffness in murine aorta. (**A**) Circumferential stress–stretch curves of WT (blue) vs eNOS^−/−^ (green) male mice carotid arteries (11 weeks male mice; WT=3, eNOS^−/−^=5) as determined by pressure myography. Black symbols indicate the results obtained at the physiological pressure range of 80–120mm Hg. Statistical differences in depends on the pressure, p-values has been calculated on stress parameters. The largest p-value is shown for simplicity of presentation. **(B)** Pressure myography analysis of glyoxal treated *ex vivo* murine arteries. Circumferential stress-stretch curves of 50mM glyoxal (red; n=6), 500μM glyoxal (green; n=4) compared to the saline one (blue; n=5) (male mice carotid arteries 11 weeks male mice). Statistical differences depend on the pressure, p-values has been calculated on stress parameters. Black symbols indicate the results obtained at the physiological pressure range of 80–120mm Hg. All data were analyzed by a two-way ANOVA with Dunnett’s post hoc analysis. p< 0,05 *; p≤ 0,001 ***. **(C)** ECM analysis of 11 weeks WT mouse carotids subjected to *in vitro* glycation treatment. TPEF carotid imaging analysis has been performed for elastin fibers (green) on saline (*a*), 500μM glyoxal (*b*), 50mM glyoxal (*c*) treated carotids; lower row panels are magnifications of the corresponding upper row panels (*a’-b’-c’*). Verhoeff-van Gieson staining has been done on saline (*d*), 500μM glyoxal (*e*), 50mM glyoxal (*f*) treated carotids. **(D)** IHC analysis of CML and AGE tissue accumulation on *in-vitro* glycated carotid murine sections. First row represents CML staining on saline (*a*), 500μM glyoxal (*b*), 50mM glyoxal (*c*); second row represent AGE staining on saline (*d*), 500μM glyoxal (*e*), 50mM glyoxal (*f*). Images have been acquired with a 20X objective.

### RAGE antagonist peptide treatment prevents aortic dilation and AGE accumulation in the aortic wall of eNOS ^−/−^ mice

We sought to determine whether the same anti-glycation strategy that mitigate endothelial function would also prevent AGE accumulation in the aorta of eNOS^−/−^ mice and mitigate ascending aortopathies. To analyze the role of RAP *in vivo* we used our established Angiotensin II osmotic pump implant model in eNOS^−/−^ mice. Angiotensin II (Ang II) chronic infusion has been previously described as inducer of increase of TAA in murine models^11^. Seven-week-old mice, fed with a hypercholesterolemic diet, were randomized in four treatment groups (Saline, Ang II, RAP (20 mg/kg), Ang II+ RAP) and infused for 28 days with Ang II (1000 ng/kg/min) or saline solution (control). Additional, mice were injected with or without RAP treatment for 28 days. Aortic aneurysms were observed in the all Ang II-treated group when compared to the control group. All animals survived until day 28. Under Ang II infusion, luminal expansion was observed throughout the ascending and descending aorta. As expected, Ang II-infused mice showed increased luminal diameter compared to control animals (1.27±0.02 vs. 1.51±0.06 mm) (**Figure 6A**). ROS accumulation was assessed as a positive control of aortic remodeling accordingly to our previous works^11^. Interestingly, our echocardiogram results indicate that RAP treatment prevents aortic dilation caused by Ang II (Ang II+RAP, 1.34±0.04 mm)(**Figure 6B**). Moreover, we observed CML and RAGE staining accumulation in the aorta of Ang II-treated mice compared to the saline animal (**Figure 6C, *a-a’* and *b-b’***). Notably, the simultaneous Ang II and RAP treatment appears to preserve from CML, glucosepane, HMGB1, and RAGE increased accumulation in the tissue (**Figure 6C, *a’’* and *b’’***). To assess the impact of RAP on AGE-mediated aortic remodeling, we performed an EVG staining on eNOS^−/−^ aortas. Ang II induced a significant remodeling of the elastin fibers compared to control. RAP treatment prevented the elastin fragmentation induced by Ang II (**Figure 6D**). These data suggested that RAGE antagonist peptide treatment is preventing AGEs accumulation in the aortic wall and is preventing RAGE pathway accumulation.

**Figure 6:**
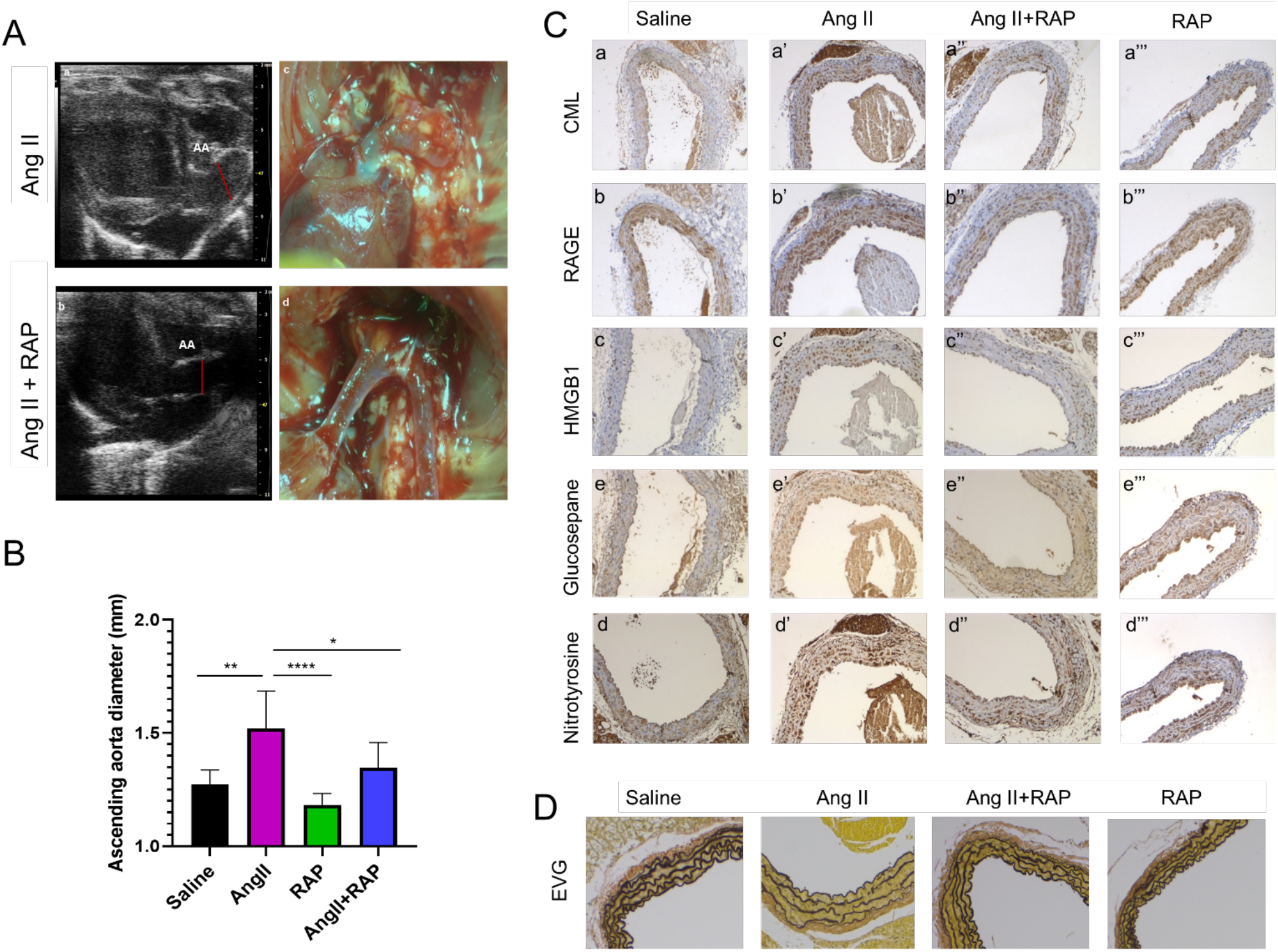
RAGE antagonist peptide prevents aortic dilation and AGEs accumulation in the aortic wall of eNOS −/−. **(A)** Upper panel: Left pictures are representative images of echocardiography analysis of Ang II and Ang II+RAP treated mice. Right pictures are representative images of ascending aorta in the mouse thoracic cavity. Lower graph: Quantification of the ascending aorta diameters (mm) acquired during the echocardiography analysis. eNOS^−/−^ mice Saline (n=6), AngII (n=11), RAP (n=8), AngII+RAP (n=6). Results are the average ± SD of ≥ 3 independent experiments. Results were analyzed by a one-way ANOVA with Dunnett’s post hoc analysis.; p< 0,05 *; p≤ 0,01 **; p≤ 0,001 ***. **(B)** IHC analysis of AGE markers (CML, RAGE, HMGB1), ROS markers (Nitrotyrosine) and crosslinker (glucosepane) tissue accumulation in eNOS^−/−^ treated mice aortic sections. **(C)** Elastin EVG staining on eNOS^−/−^ treated mice aortic sections.

## DISCUSSION

Thoracic aortic aneurysm is characterized by a significant increase of the artery diameter due to an intrinsic weakness of the aortic wall.^34^ TAA rarely manifest with symptoms and for has been defined as a “ silent killer”^35, 36^. Indeed, as 95% of the patients are asymptomatic, they can easily guide to dreadful complications^37^. About 22% of people die before reaching the hospital because of an aortic dissection or rupture ^38^. More knowledge has been accumulated on the natural etiology of TAAs in terms of size at rupture or dissection, expansion rates and sex related outcomes. However, noninvasive imaging techniques that allow clinicians to select patients as candidates for repair based on size and symptoms are the main tools^35^. Beyond the absolute size of the aneurysm, there is a need for useful new diagnostic tools capable of detecting TAA in asymptomatic patients and that can predict aortic dissection or rupture before they happen. Our study unveils a new mechanism of AGE accumulation in the vascular wall and tests RAP as a tool to reduce both AGE-mediated endothelial hyper-permeability and AGE-mediated vascular remodeling.

In this study, we show that RAP can mitigate AGE-mediated endothelial hyper-permeability *in vitro* and impact ascending aortopathy *in vivo*. Cell culture experiments with endothelial cell contraction and migration show an effect of glyoxal of EC physiology. A Z-Theta ECIS system is used to measure endothelial hyper-permeability of EC under glyoxal stimulation with results showing that RAP rescues its effect. If *in situ* glycation is the result of infiltration of preformed glycation products and/or reactive glycation precursors into the aortic wall via impaired endothelial barrier we should expect accumulation of glycation products in the vascular wall near the endothelial barrier. Indeed, histological examination of human resected aortas from dilated and not-dilated aorta shows accumulation of generalized AGE products around the aortic lumen, and in the adventitia layer. As described in the literature, in larger vessels diffusion of solutes to the media is supplemented by vasa vasorum^21 20^. The nutrients and compounds have been shown to permeate as much as half or two-thirds of the media and that is also consistent with the results in a large majority of our specimens. Notably glucosepane, the most abundant AGE crosslinker, also accumulates in the aortic wall. This finding suggests that glycation may affect aortic wall stability via crosslinking of large ECM molecules. To assess the role of glycation on aortic wall stability we developed an *in vitro* glycation reaction that resulted in the accumulation of CML and in elastin degradation leading to ECM remodeling. To test whether impaired endothelial function is associated with altered vascular stiffness, we used a murine model with a known severe vascular phenotype characterized by hypertension and increased endothelial permeability (eNOS^−/−^). The arteries of these mice are indeed stiffer than WT ones by a pressure myography assay that measures biaxial inflation-extension tests. Since AGE could impact both the ECM directly or indirectly via VSMC phenotypic switch, we also assessed the phenotype of VSMC exposed to glyoxal with results showing that AGE decreases the contractile phenotype marker levels, including caldesmon, calponin, myocardin, SMMHC and SMA and, increases the synthetic ones like vimentin, CTGF. We then tested whether an anti-glycation strategy, could prevent AGE accumulation in the aorta of either eNOS^−/−^ mice with data showing that RAP improved the architecture of the ascending aorta and reduced CML, RAGE, HMGB1 and glucosepane accumulation along with reducing ascending aortic dilation by echocardiographic measurements. Finally, we validate the direct action of AGE on vascular stiffness by exposing murine carotid arteries to glyoxal *ex vivo*, with results showing increased vascular stiffness. Overall, this study provides insight into the mechanism underlying vessel stiffness and vascular dysfunction in thoracic aortic aneurysm studying the effect of AGEs on ECM remodeling and cellular component of the aortic wall.

AGE and RAGE have been described to be involved in the pathogenesis of numerous diseases including hypertension^39^, myocardial infarction^40^, and post-percutaneous coronary interventional restenosis^41^. The focus on the implication of AGEs in aortic aneurysm however has been poorly highlighted. The function of the thoracic aorta is the result of a dynamic, highly preserved micro-structural organization, which correlates with both hemodynamic conditions, as well as the cellular and extracellular components of the aortic wall. Growing evidence suggests that altered wall stress distribution and predisposing genetic background contribute to the risk of develop aortopathies^42^. Dissection areas are also more prone to lead to endothelial dysfunction, thus the role of glycation in mediating ascending aortic remodeling could also be impacted by altered hydrodynamic condition that impact endothelial cell physiology independently of AGE. Our data demonstrate that aortic ECs show a decrease of VE-cadherin expression and an increase of barrier permeability induced by glyoxal. Altered shear stress and mechanotransduction are also known regulators of cell-cell adhesion, however the clinical data on the frequency of aortic dissection in T2DM patients corroborate the idea that accumulation of glycation products is linked to endothelial dysfunction and vascular stiffness. Glycation strongly contributes to structural and functional degeneration of tissues and diseases directly modifying the extracellular matrix proteins via crosslinking and modification of protein interactions, or modulating the cellular and inducing an inflammatory reaction via glycation product-mediated receptor signaling^43^. Cross linking caused by AGEs leads to vascular stiffening and endothelial dysfunction ^44^, but the mechanism through which glyoxal causes vascular dysfunction remains still unclear.

Several limitations are associated with this study: first, validation should be obtained using patient-derived endothelial cells from dilated and not-dilated ascending aorta. We decided to work with commercially available EC to avoid the role of patient-comorbidities in this first study as this will require assessing a large number of patients which are not yet available in our lab. Another limiting factor is the use of glyoxal as a precursor of glycation products. Protein modifications could include early glycation products (EGPs) as well as advanced glycation end-products (AGEs). Early glycation involves attachment of glucose on ε-NH_2_ of lysine residues of proteins leading to generation of the Amadori product (an early glycation species)^87^. Thus, Amadori product will be tested in future studies. Glycation precursors will be generated by incubation with the various glycation precursors and purified by chromatographic filtration. Alternative mechanisms to consider are the reported role of AGEs on DNA damage and that our pressure myograph system has an intrinsic limitation of vessel diameters that forced us to study descending aorta and carotid arteries instead of ascending aortas. In conclusion, this work provides strong evidence that glycation affects aortic wall stability impacting VSMCs and endothelial cell physiology. The result leads to compromised ECM structure by inducing stiffness of the tissue that potentially increases the risk of developing aortopathies. Metric measurements are imperfect predictors of aortic dissection and rupture as they do not adequately assess aortic wall remodeling and stress, thus this study is the first step into a mechanistic approach to explain aortic remodeling independently of ascending aortic diameter.

## MATERIALS AND METHODS

### Patient selection and surgical procedures

According to the approved IRB protocol, a retrospective study was performed on patients enrolled at: Columbia University (IRB # AAAR6796), The Valley Cardiac Biobank (IRB # 20132309), University of Pennsylvania (IRB # 809349). All patients included in this study have been followed for aortic valve diseases (stenosis or insufficiency) and/or enlargement of the ascending aorta and reached the criteria for surgical intervention. All patients provided written informed consent. Blood was taken before surgery and patients were divided in two groups according to the morphology of the aortic valve assessed by transesophageal echocardiography (TEE), computed tomography (CT scan) or both, and confirmed by intraoperative assessment. Exclusion criteria were: genetic disease associated with TAA, Bicuspid Aortic Valve Syndrome, connective tissue disease, chronic inflammatory disease, previous myocardial infarction, severe heart failure (NYHA III, IV), endocarditis and active cancer. Non-dilated patients presented with a mean aortic diameter of 4.59±0.89 cm. Dilated patients presented with a mean aortic dilatation equal to 6.07±0.51 cm. Of the 30 patients who match the inclusion criteria, 15 had ascending aortic repair with aortic valve replacement, 11 had ascending aortic repair and 4 underwent aortic valve replacement (**Table S1**).

### Cell culture

Primary Aortic Smooth Muscle Cells, Normal, Human (HASMC) (Cat# PCS-100-12. ATCC); Bovine Aortic Endothelial cells (BAOEC) (Cat# B304-05. CELL Applications, Inc.). HASMCs and BAOECs were grown in DMEM medium completed with glutamine, penicillin/streptomycin solution, and 10% FBS (Corning).

### Antibodies and reagents

Glyoxal solution (Cat# 50649) was from Sigma-Aldrich. Rabbit anti-VE-Cadherin (#2158) was from Cell Signaling and was diluted 1:1000 for Western blot analyses. Anti-VE-Cadherin antibody was diluted 1:100 for immunofluorescence assay. Horseradish peroxidase (HRP) conjugated goat anti-rabbit (sc-2054) secondary Ab was from Santa Cruz Biotechnology, while HRP-goat anti-mouse (115-035-003) was from Jackson ImmunoResearch Laboratories. Alexa Fluor 488 donkey anti-rabbit (A21206), Alexa Fluor 555 Phalloidin (A34055) and DAPI (D3571) were from Thermo Fisher Scientific. GAPDH (G-9) monoclonal HRP has been used in western blot 1:5000 Santa Cruz Biotechnology. RAGE Antagonist, FPS-ZM1 – Calbiochem was from Sigma-Aldrich.

### Western Blot

Cultured BAOECs were treated with RAP [5μg/ml] for 2h and then glyoxal 500μM was added to the medium for 24h. ECs were lysed in RIPA lysis buffer (Boston BioProducts) supplemented with Phospatase inhibitor (Thermo Fisher Scientific) and EDTA-free Protease Inhibitor Cocktail (Roche). Cellular lysates were incubated for 20 min on ice, and then centrifuged at 15,000 g, 20 min, at 4 °C. The total protein amount was determined using the bicinchoninic acid (BCA) protein assay reagent (Thermo Fisher Scientific). Specific amounts of protein were separated by SDS–PAGE with precast Bolt 4-12% Bis-Tris gel (Thermo Fisher Scientific) or Mini PROTEAN TGX precast 4-20% (Bio-Rad). Proteins were then transferred to a nitrocellulose membrane (Bio-Rad), probed with Abs of interest and detected by enhanced chemiluminescence technique (Pierce_ Thermo Fisher Scientific).

### Immunohistochemistry

Human aortic tissues, mouse aortas and carotids tissues were fixed in 10% formalin for 24hrs, transferred to EtOH 70%. Embedding in paraffin and sectioning (5μm-thick) of the samples have been performed by the histological core facility. Antigen retrieval was performed by an incubation in 1x sodium citrate buffer (Sigma Aldrich) at 100°C for 30’. Samples were saturated in a blocking solution of 3% BSA (Boston BioProduct) 1xPBS for 1h RT. Anti-AGE (Abcam, Cambridge, UK #ab23722, 1:1000), anti-CML (Abcam #ab27684, 1:500), anti-RAGE (Abcam #ab3611, 1:1000) and anti-glucosepane (David Spiegel laboratory, Yale University, New Haven, CT, per material transfer agreement, 1:100 dilution) antibodies were diluted in DAKO primary antibody diluent (Agilent Technologies, Santa Clara, CA, USA) and applied to sections overnight at 4°C. DAKO HRP polymer-conjugated anti-rabbit and anti-mouse secondary antibodies (Agilent, #ab214880, #ab214879) were applied at room temperature for 1hr. Washes were accomplished using 1x dilution of DAKO Wash Buffer 10x (Agilent). IHC stains were developed for 6 minutes using 3,3’-diaminobenzidine tetrahydrochloride substrate (Abcam) and sections were counterstained for 3 minutes with Gill’s hematoxylin (SigmaAldrich, St. Louis, MO, USA). Slides were mounted using Perimount mounting medium (Fisher Scientific, Hampton, NH, USA) and imaged by a Leica DMi1 Inverted Phase Contrast Microscope with a 5X or 10X objectives (Leica Camera AG, Wetzlar, DE). Histological analysis of the human aortic tissues was performed by Movat Pentachrome staining for proteoglycans and collagen, and Verhoeff-Van Gieson staining for elastin according to the protocol of the Histology Laboratory of the University of Pennsylvania School of Medicine and the Molecular Pathology/MPSR (HICCC) histology core facility at Columbia University.

### Endothelial Permeability assay

Cell barrier function was monitored with ECIS Z-theta instrumentation from Applied BioPhysics. ECIS culture plate 8W10E+ was coated with gelatin for 15’ at RT. 65000/well ECs were plated in each well, except for the control free cell wells, in DMEM 10% FBS medium. The plate is left at RT for 15’. The plate is then put in the machine at 37°C and the assay start. The resistance (R, ohm), impedance (Z, ohm), capacitance (C, nF) values are monitored over time. In around 48h, the R value, given by the cell monolayer, reached the plateau. Change the medium and wait for the plateau to be reached again in few hours. Cells are, then, pretreated with RAP [5μg/ml] or [10μg/ml] for 2h and then glyoxal 500μM was added to the medium. The R and C values are monitored over time.

### Cell viability assay

Cell viability was evaluated by CellTiter-Glo Luminescent assay (Cat# G7572. Promega). Briefly, an equal number of cells (HASMC and BAOEC 20000/well) were seeded in 96-well culture plates in a 100μl volume of complete medium. Cells were treated with various concentrations of glyoxal (50mM, 10mM, 5mM, 1mM, 500μM) in complete medium and control wells (PBS) were also prepared. The assay plate was equilibrated at room temperature for 30’. The CellTiter-Glo reagent was prepared according to the manufacturer protocols and an equal volume of reagent was added to the volume of the cell culture medium present in each well (100μl). A blank point (no cells, no medium. 200μl of CellTiter-Glo reagent) was used as an internal control of the plate. The plate was placed on an orbital shaker for 2’ to induce the lysis of the cells. Then the plate was incubated at room temperature for 10’ to stabilize luminescent signal. The 200μl mix volume was transferred to an opaque-wall 96 well plate and the luminescence (RLU) was red by SpectraMax iD3 Multi-Mode Microplate Readers (Molecular Devices). All the experimental points of the assay were prepared in triplicate. The average value of the blank wells was subtracted to each experimental point of the assay. Luminescence values are calculated relative to control. Values represent the mean ± S.D. of X replicates for each cell number.

### Three-dimensional collagen gel assay

The *in vitro* contraction assay has been performed according to the protocol of the Cell Contraction Assays, Floating Matrix Model (Cat# CBA5020. Cell Biolabs) kit. Briefly, 3 × 10^6^/ml HASMCs, BAOEC were mixed with a collagen solution and seeded in the 24 well plate provided by the kit. After 1h at 37°C, complete medium +/− glyoxal treatment (1mM glyoxal for HASMC, 500μM glyoxal for ECs) was added to the polymerized collagen gel. The collagen gel size changing (contraction index) was monitored over time and pictures were taken after 24h. Analysis of the results was performed using the ImageJ software package. The contraction index calculation derive from the formula: (well area - gel area)/(well area).

### Immunofluorescence and Confocal microscopy on cultured ECs

Cells were plated on 0.17 mm glass coverslips (no. 1.5) pre-coated with gelatin and allowed to adhere to a monolayer. BAOECs were pre-treated with RAP [5μg/ml] for 2h and then glyoxal 500μM was added to the medium for 24h. Cells were washed in phosphate-buffered saline (PBS), fixed in 4% para-formaldehyde (PFA), permeabilized in 0.1% Triton PBS 1X for 2 min on ice, incubated with different primary Abs and revealed by appropriate Alexa-Fluor-tagged secondary Abs (Thermo Fisher Scientific). Cells were analyzed by using a Nikon Ti Eclipse inverted microscope with A1 scanning confocal microscope at the Confocal and Specialized Microscopy Shared Resource (CSMSR). 60x/1.49 Apo TIRF Oil immersion objective was employed. 1024×1024 pixel images were acquired using NIS Elements software. The acquisition was performed by adopting a laser power, gain, and offset settings that allowed maintaining pixel intensities (gray scale) within the 0– 255 range and hence avoid saturation. Analysis of confocal images was performed using the ImageJ software package.

### RNA isolation and real-time PCR assay

HASMCs were seeded in a 6xwell plate at a confluence of 250000 cells/well in DMEM 10%FBS. The day after cells were treated with 1mM glyoxal in DMEM 10% FBS at 37°C for 24h. After 24h of treatment, cells were washed three times with PBS and frozen at −80°C. RNA isolation and reverse transcription: total RNA was extracted following the manufacturer’s recommended protocol (RNeasy Mini Kit, Qiagen). The quality and integrity of the total RNA were quantified by the NanoDrop DeNovix. cDNAs were generated using the High Capacity cDNA Reverse Transcription Kit (Applied Biosystems). TaqMan real-time RT-PCR assay (Thermo Fisher Scientific): mRNA expression of *SMA* (Hs00426835_g1), *SMMHC* (Hs00975796_m1), *MYOC* (Hs00538071_m1), *VIMENTIN* (Hs00958111_m1), *CTGF* (Hs00170014_m1), *Coll1A1* (Hs00164004_m1) and the endogenous housekeeping control gene, *GAPDH* (Hs99999905_m1). Sybr Green real-time RT-PCR assay probes (Roche Universal Probe library): Fw 5’gagcgtcgcagagaacttaga3’ and Re 5’ ttcctctggtaggcgattctt3’ (*CALDESMON*), Fw 5’ gctgtcagccgaggttaaga3’ and Re 5’ cctcgatccactctctcagc3’ (*CALPONIN*), Fw 5’agccacatcgctcagacac3’ and Re 5’ gcccaatacgaccaaatcc3’ (*GAPDH*). mRNA expression was measured on a QuantStudio 7 Flex Real-Time PCR System (Applied Biosystems). For each sample, three technical replicates of each gene were run in a 384-well plate (cDNA concentration 50 ng/well) according to the manufacturer’s protocol. The experimental threshold (Ct) was calculated using the algorithm provided by the Applied Biosystems qPCR analysis software. The experimental Ct values were converted into relative quantities using the ΔCt method described by Pfaffl and amplification efficiency of each gene was calculated using a dilution curve and the slope calculation method (Pfaffl, 2001). We chose as normalization factor GAPDH values.

### Wound healing assay

250000 ECs and 200000 HASMCs were plated in each well of a 24-well plate and incubated overnight at 37 °C, 5% CO2 in a humidified atmosphere. Once the monolayer of cells has been reached, cells were starved for 2h in 0.5% medium. Then each well was scratched with a 200μl pipette tip in the middle of the monolayer. Cells were then washed once with PBS and treated with 1mM (HASMC) and 500μM (BAEC) glyoxal or PBS in complete medium for 24h at 37 °C. The assay was imaged by a Leica DMi1 Inverted Phase Contrast Microscope with a 5X objective. Images of the wound closure were taken at 0h (T0), 6h and 24h (T24). Displacement of the cell sheet leading edge over time was quantified by Image J.

### Mouse strain and maintenance

Animals were treated according to the guidelines and under the supervision of the Institute of Comparative Medicine (ICM) at Columbia University. C57BL/6 wild-type (WT) and eNOS null mice (B6.129P2-Nos3) on the C57BL/6 background were obtained from Jackson Laboratories (Bar Harbor, ME) and euthanized by isoflurane overdose (5% isoflurane inhalation), thoracotomy and exsanguination from the heart (cardiac puncture).

### Mouse osmotic pump implantation

Angiotensin II chronic infusion model. 7 weeks old wild type male mice (C57BL/6J) were fed with hypercolestermomic diet and infused with saline or Angiotensin II (1000 ng/kg/min) using osmotic pumps (Alzet 2004) for 28 days as previously described. Moreover, mice have been injected with or without RAP treatment (FPS-ZM1 20 mg/kg, Sigma-Aldrich) for 28 days. X treated and Y untreated mice were used. Mice were sacrificed according to the IACUC protocol. Tissues like carotids, ascending and descending aortas were harvested after 28 days. Experiments were conducted under the approved IACUC protocol AC-AAAU6474. At baseline and 28 days after aneurysm induction, aortic expansion was measured using echocardiographic evaluation (Oncology Precision Therapeutics and Imaging Core/OPTIC (HICCC) Columbia University). Cross-sectional analysis was performed by experienced sonographers using a Visulasonics VEVO 3100 High Frequency Ultrasound Imaging System.

### Pressure myography assay

Vessels were harvested from 11 weeks old male eNOS^−/−^ or WT mice. After animal sacrifice, one or both carotid arteries were isolated, cleaned of fat (except at the two ends; see vessel mounting procedure below), stored at −80°C in PBS, and then thawed for 30’ and warmed to 37°C for 30’ prior to myograph testing. We chose to analyze mouse carotid arteries, because are relatively straight vessels of reasonable length (2.5-4.2 mm) and are lacking branches. The carotid was mounted on 300μm stainless steel – P/PP/CP grooved cannulae and secured using a 7-0 silk suture with two square knots on each side of the vessel. A DMT model 114P Pressure Myograph was used to determine the stiffness of the vessels. Measurements were continuously recorded with dimension analysis MyoVIEW software. The experimental protocol applied in the assay is adapted from the method described by Brankovic and colleagues ^16^.

### Ex vivo glycation of murine vessels

The entire thoracic aorta connected to the left and right carotids were exposed in a WT mouse. Murine carotid from WT animals was exposed, cleaned of fat and connective tissue, and cut on one edge. After rinsing with PBS the inner side of the carotid artery, one edge of the vessel was clamped with 7-0 silk suture and infused with a glycation treatment (glyoxal 50mM or 500μM) or saline solution in DMEM 10% FBS. The second edge of the vessel was also clamped, after the injection of the treatment, to create a double clipped-artery and the tissue was incubated at 37°C for 24h. A DMT model 114P Pressure Myograph was used to determine the stiffness of the vessels. Measurements were continuously recorded with dimension analysis MyoVIEW software.

### Second harmonic generation (SHG) imaging

Paraffinized tissues were deparaffinized using the same protocol as described in IHC and mounted with Permount. SHG and TPEF imaging were performed on a Nikon A1RMP multiphoton confocal microscope (Nikon; Minato, Tokyo, Japan). The SHG and TPEF signals were generated using either 860-nm (for SHG) or 700-nm (for TPEF) excitation light from a Ti:sapphire femtosecond laser and detected using Nikon Apo LWD 25x/NA1.1 gel immersion as an objective with a 400-450 nm bandpass filter. All images were acquired at 1024×1024 resolution using NIS Elements software.

### Statistical analysis

For *in vitro* assays data are presented as the means ± SD in this study. Prism/Excel software was used to analyze the data. One-way analysis of variance (ANOVA) and Student’ t-tests were used to analyze the differences between groups. Differences were considered statistically significant when p value is < 0.05. For pressure myograph Diameter–tension relationships were analyzed by calculating the linear regression lines and their corresponding slopes and diameter at force 0 (Y0), which were then compared among groups using the Student t test or one-way ANOVA. Diameter–pressure curves were compared with two-way ANOVA followed by the Sidak multiple comparisons tests. Stress–strain curves were analyzed by extracting the β value for each curve which we compared among groups using the Student t test. Aortic cross-sectional area curves obtained by MRI and collagen immunofluorescence measurements were compared by the Mann–Whitney U test. Results were considered statistically significant at p-values <0.05. Statistical analysis was performed with GraphPad Prism 7 software or R.

## ACKNOWLEDGMENTS

Funding: This work was supported by National Institutes of Health R01s HL122805 and HL143008 (GF) and T32-HL007343 (AF), T32HL007854 (AA), AATS Norman E. Shumway Research Scholarship (DL), The Kibel Fund for Aortic Valve Research (to GF), The Valley Hospital Foundation ‘Marjorie C Bunnel’ charitable fund (GF and JBG) and Andrew Sabin Family Foundation Cardiovascular Research Laboratory (GF). The Confocal and Specialized Microscopy Shared Resource, the Oncology Precision Therapeutics and Imaging Core (OPTIC) and the Molecular Pathology core at the Herbert Irving Comprehensive Cancer Center are supported by NIH grant #P30 CA013696 (National Cancer Institute).

The authors declare no conflict of interest.

## Notes

### Competing Interest Statement

The authors have declared no competing interest.

