## Supplemental Material for "RAGE antagonist peptide mitigates AGE-mediated endothelial hyperpermeability and accumulation of glycoxidation products in human ascending aortas and in a murine model of aortic aneurysm"

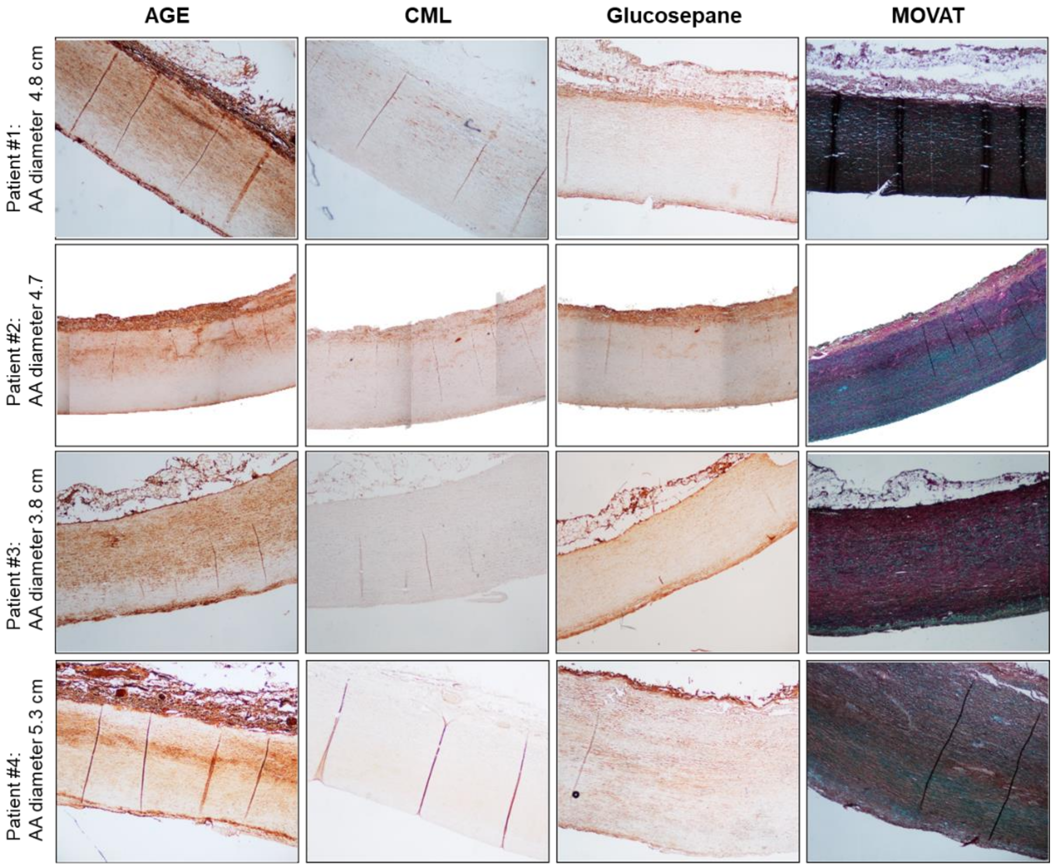

**Figure S1**: **AGEs accumulate on the intima and adventitia layer of the aorta, independently of the ascending diameter.** Representative images of patient tissue sections selected from our cohort. Human ascending aorta cross-section stained for general AGE, CML, glucosepane and Movat. Patients present a similar pattern of staining regardless the diameter of the ascending aorta, sex, and key co-morbidities.

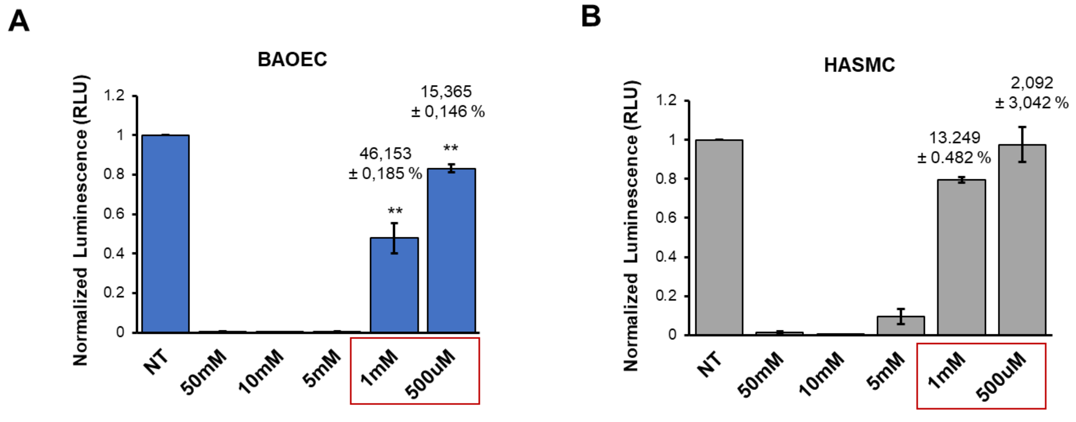

**Figure S2: Cell viability assay after glyoxal treatment on ECs and VSMCs.** Cell viability was evaluated by CellTiter-Glo Luminescent assay. Cell viability assay has been performed on cells treated or not with several glyoxal concentrations (50mM, 10mM, 5mM, 1mM, 500µM) for 24h. 50mM has been used as a negative control. Values are expressed as normalized luminescence relative to the NT cells. **(A)** BAOECs viability is decreased by 46.153 ± 0.185% and 15.365 ± 0.146 % with 1mM and 500µM glyoxal treatment respectively. **(B)** HASMCs viability is decreased by 13.249 ± 0.482% and 2.092 ± 3.042 % with 1mM and 500µM glyoxal treatment respectively. Results are the average ± SEM of ≥ 3 independent experiments. Results were analyzed by a two-tailed heteroscedastic Student's t-test; $p< 0,05$ *; $p\leq 0,01$ **.

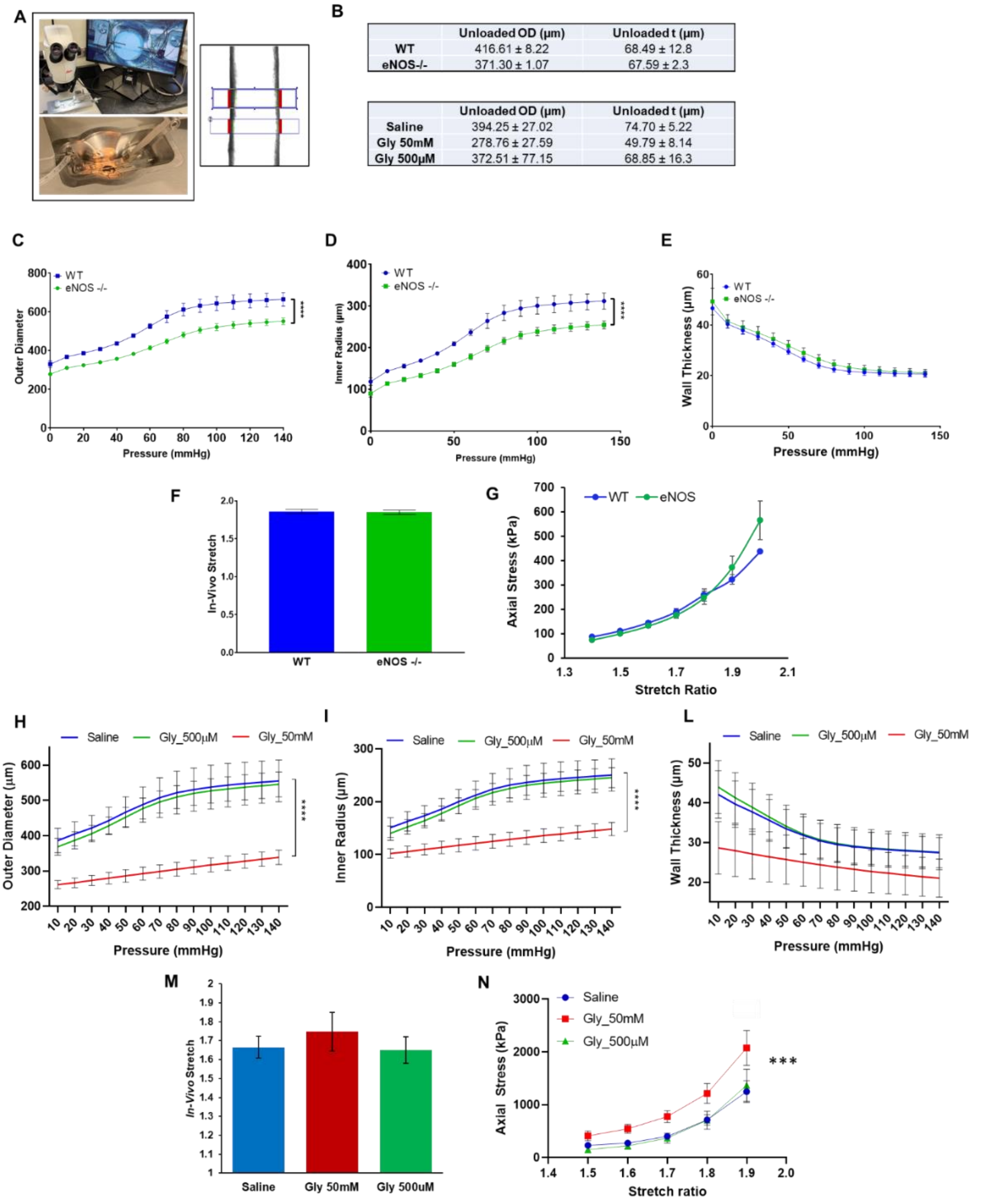

**Figure S3: (A)** Explanted carotid from an 11-week-old mouse loaded in the pressure myograph system. The outer diameter measurements are captured from two independent boxes designed on the vessel (red lines). Images of the DMT model 114P Pressure Myograph and of the cannulae on which the vessel is mounted. **(B)** Unloaded outer diameter and thickness dimensions of carotid arteries from WT and eNOS$^{-/-}$ mice and from saline and glyoxal treated vessels. Vessel dimensions of WT control and eNOS-null mice. Loaded vessel outer diameter **(C)**, inner (lumen) radius **(D)**, and wall thickness **(E)** of WT (n=3) and eNOS-null (n=5) right carotid arteries. Data show mean ± SEM. All data were analyzed by ANOVA. **(F)** Mean In-Vitro Stretch (IVS) measurements of both groups. **(G)** Axial stress-stretch curves of the WT and eNOS carotid arteries were determined at 100 mm Hg. Results in all panels show mean ± SEM. All data were analyzed by ANOVA. Data show mean ± SEM. All data were analyzed by a two-way ANOVA with Sidak's post hoc analysis. **(H)** Loaded vessel outer diameter, **(I)** inner (lumen) radius and **(L)** wall thickness of saline (blue; n=5), 500µM glyoxal (green; n=4), 50mM glyoxal (red; n=6) carotid arteries. **(M)** Mean In-Vitro Stretch (IVS) measurements of the 3 groups. **(N)** Axial stress-stretch curves of saline, gly500µM and gly50mM treated carotid arteries were determined at 100 mm Hg. Data show mean ± SEM. All data were analyzed by ANOVA. p< 0,05 *; p≤ 0,01 **; p≤ 0,001 ***.

| | Overall | Dilated | Non-dilated |
|---|---|---|---|
| | (n=30) | (n=9) | (n=21) |
| **Age (mean (SD))** | 64.11 (±10.20) | 63.22 (±12.02) | 64.53 (±9.55) |
| **Sex** | | | |
| **Male** | 21 (75.0%) | 7 (77.8%) | 14 (66.7%) |
| **Female** | 9 (25.0%) | 2 (22.2%) | 7 (33.3%) |
| **Hypertension** | 20 (66.7%) | 5 (55.6%) | 15 (71.4%) |
| **Hyperlipidemia** | 15 (50.0%) | 4 (44.4%) | 11 (52.4%) |
| **Diabetes Mellitus** | 3 (10.0%) | 1 (11.1%) | 2 (9.5%) |
| **Smoking** | 16 (53.3%) | 4 (44.4%) | 12 (57.1%) |
| **Coronary Artery Disease** | 9 (30.0%) | 4 (44.4%) | 5 (23.8%) |
| **Coronary Artery Bypass Graft** | 2 (6.9%) | 0 (0.0%) | 2 (10.0%) |
| **Aorta Diameter (mean (SD))** | 5.03 (±1.04) | 6.07 (±0.51) | 4.59 (±0.89) |
| **Surgery Performed (%)** | | | |
| **AA** | 11 (36.7%) | 4 (44.4%) | 7 (33.3%) |
| **AVR** | 4 (13.3%) | 0 (0.0%) | 4 (19.0%) |
| **AVR/AA** | 15 (50.0%) | 5 (55.6%) | 10 (47.6%) |
| **Tricuspid Aortic Valve (%)** | 30 (100.0%) | 9 (100.0%) | 21 (100.0%) |

**Table S1: Patient demographic and characteristics**
